# Comparative exometabolome and extracellular-vesicle metabolome of *Bifidobacterium*

**DOI:** 10.64898/2026.11.18.767755

**Authors:** Eveliina Munukka, Mafiur Rahman, Katja H. Salonen, Ifrat Tamanna, Nasir Badal, Mohd Umar Tabrez, Marianne Gunell, Teemu Kallonen, Matej Orešič, Alex M. Dickens, Antti Hakanen, Pieter C. Dorrestein, Md. Khirul Islam, Santosh Lamichhane

## Abstract

*Bifidobacterium* species are early colonisers of the human gut and are widely used as probiotics, yet their extracellular chemistry remains largely unexplored, and the small-molecule cargo of their extracellular vesicles (EVs) has not been characterised. We carried out an exploratory, descriptive characterisation of the exometabolome and EV metabolome of *Bifidobacterium longum* 35624 and *Bifidobacterium animalis* subsp. *lactis* at 24 and 48 h of anaerobic growth, using untargeted LC–MS/MS coupled with feature-based molecular networking and semi-targeted analysis of conjugated lipids. The two strains reshaped the shared culture medium through distinct, strain-specific nutrient conversions. *B. longum* 35624 deamidated nicotinamide to nicotinic acid and produced the tryptophan catabolite indole-3-lactic acid, whereas *B. animalis* subsp. *lactis* catabolised purine nucleosides and accumulated hypoxanthine; the two strains also differed in carbohydrate utilisation. EVs, purified by ultracentrifugation and size-exclusion chromatography and characterised by nanoparticle tracking analysis, immunoassay and electron microscopy, carried a sparse small-molecule cargo. A membrane glycolipid represented the clearest vesicle-associated signal of bacterial origin, whereas bile acid and *N*-acyl lipid conjugates were also detected but could not be attributed to bacterial production, as they were present in the culture medium. Together, these data provide a first reference description of the *Bifidobacterium* exometabolome and EV metabolome and identify strain-specific extracellular metabolic activities as priorities for targeted validation.

## Introduction

The human gut microbiome is a metabolically active organ whose biochemical repertoire extends far beyond the capacity of the host genome. Using dietary, environmental and host-derived substrates, intestinal bacteria transform and generate a structurally diverse set of small molecules that regulate epithelial function, immune homeostasis, bile acid metabolism, xenobiotic biotransformation and neuroendocrine signalling. Microbiome-derived metabolites are principal mediators of host–microbe interaction, providing a functional link between microbial ecology and human physiology^1,2^. Advances in high-resolution mass spectrometry, molecular networking and computational metabolomics have transformed our ability to interrogate this microbial chemistry. Moving beyond fermentation products such as short-chain fatty acids, gut bacteria synthesise modified amino acids, complex lipids, secondary bile acids and numerous conjugated molecules that remain underexplored^2^. Conjugated metabolites in particular have changed our understanding of the scope of host–microbe chemical communication. Quinn and colleagues showed that the gut microbiota synthesises previously unknown amino acid-conjugated bile acids, including phenylalanocholic, tyrosocholic and leucocholic acid, revealing a class of microbiome-derived signalling molecules capable of engaging host bile acid receptors^3^. Subsequently, our group and laboratories worldwide have expanded this repertoire to include diverse amino acid and lipid conjugates, indicating that microbial conjugation is a widespread but still poorly characterised feature of intestinal metabolism^4^.

*Bifidobacterium* species are among the earliest colonisers of the infant gut and remain members of the healthy adult microbiome throughout life^5^. Experimental and clinical studies have linked bifidobacteria to improved epithelial barrier function, immune maturation, resistance to enteric pathogens, control of intestinal inflammation and metabolic homeostasis, and strains of both *Bifidobacterium longum* and *Bifidobacterium animalis* are used clinically as probiotics^6^. Work on bifidobacterial metabolism has nevertheless focused on carbohydrate utilisation, human milk oligosaccharide degradation, acetate production, lactate metabolism and cross-feeding, leaving most of their extracellular chemical diversity unexplored^7^. The extracellular metabolome, or exometabolome, is the functional interface through which microorganisms interact with neighbouring microbes and the host. Unlike intracellular metabolites, which mainly reflect cellular metabolism, extracellular metabolites comprise secreted molecules, transformed environmental substrates, metabolic by-products and signalling compounds.

Untargeted profiling of the exometabolome therefore reports on microbial functional activity in a way that cannot be inferred from genomic or taxonomic data alone^1,3^. A further and comparatively unexplored dimension of microbial extracellular chemistry is represented by bacterial extracellular vesicles (EVs). These nanoscale membrane-derived particles are produced by both Gram-negative and Gram-positive bacteria and transport proteins, lipids, nucleic acids, polysaccharides and other bioactive molecules, enabling communication within microbial communities and between microorganisms and host tissues^8,9^. Gut commensal vesicles can themselves shape host physiology; vesicles from *Akkermansia muciniphila*, for example, modulate epithelial barrier integrity^10^, and recent work has shown that gut microbial vesicles carry defined nucleic acid and metabolic cargo^11,12^. Although the proteomic, lipidomic and genomic composition of bacterial EVs has been studied extensively, remarkably little is known about their small-molecule cargo, and almost all of the available data concern Gram-negative outer-membrane vesicles^13^. Whether vesicles selectively package conjugated microbial metabolites, rather than simply mirroring the surrounding extracellular metabolome, is unknown for probiotic bacteria such as *Bifidobacterium*.

We therefore undertook an exploratory, discovery-phase characterisation of the extracellular metabolome and the extracellular vesicle metabolome of *B. longum* 35624 and *B. animalis* subsp. *lactis*, using untargeted LC–MS/MS integrated with feature-based molecular networking and knowledge-guided annotation. By comparing culture supernatants, purified vesicles and medium controls across growth, we set out to map the extracellular chemical landscape of these organisms and to identify conjugated microbial metabolites systematically. The study is deliberately hypothesis-generating: our aim is to establish what chemistry is present and to define a tractable set of candidate molecules, rather than to test the activity of any one of them. To our knowledge, this is the first description of the small-molecule conjugated metabolome of *Bifidobacterium* extracellular vesicles.

## Results

### Experimental workflow and characterisation of bacterial extracellular vesicles

Fig. 1 shows the overall study design used to investigate the extracellular chemistry of *B. longum* 35624 and *B. animalis* subsp. *lactis*. We performed a time-series in vitro culturomics experiment in which both strains were cultivated under strictly anaerobic conditions in Fastidious Anaerobe Broth (FAB). Three independent biological replicates were prepared per strain and sampled before inoculation (uninoculated medium), immediately after inoculation, and at 24 h and 48 h of growth (Supplementary Table 1). Growth was monitored spectrophotometrically (OD_600_). Cell-free supernatants were collected for exometabolomic profiling, and parallel cultures were processed for EV isolation, allowing direct comparison between the soluble extracellular metabolome and EV-associated cargo.

**Fig. 1.**
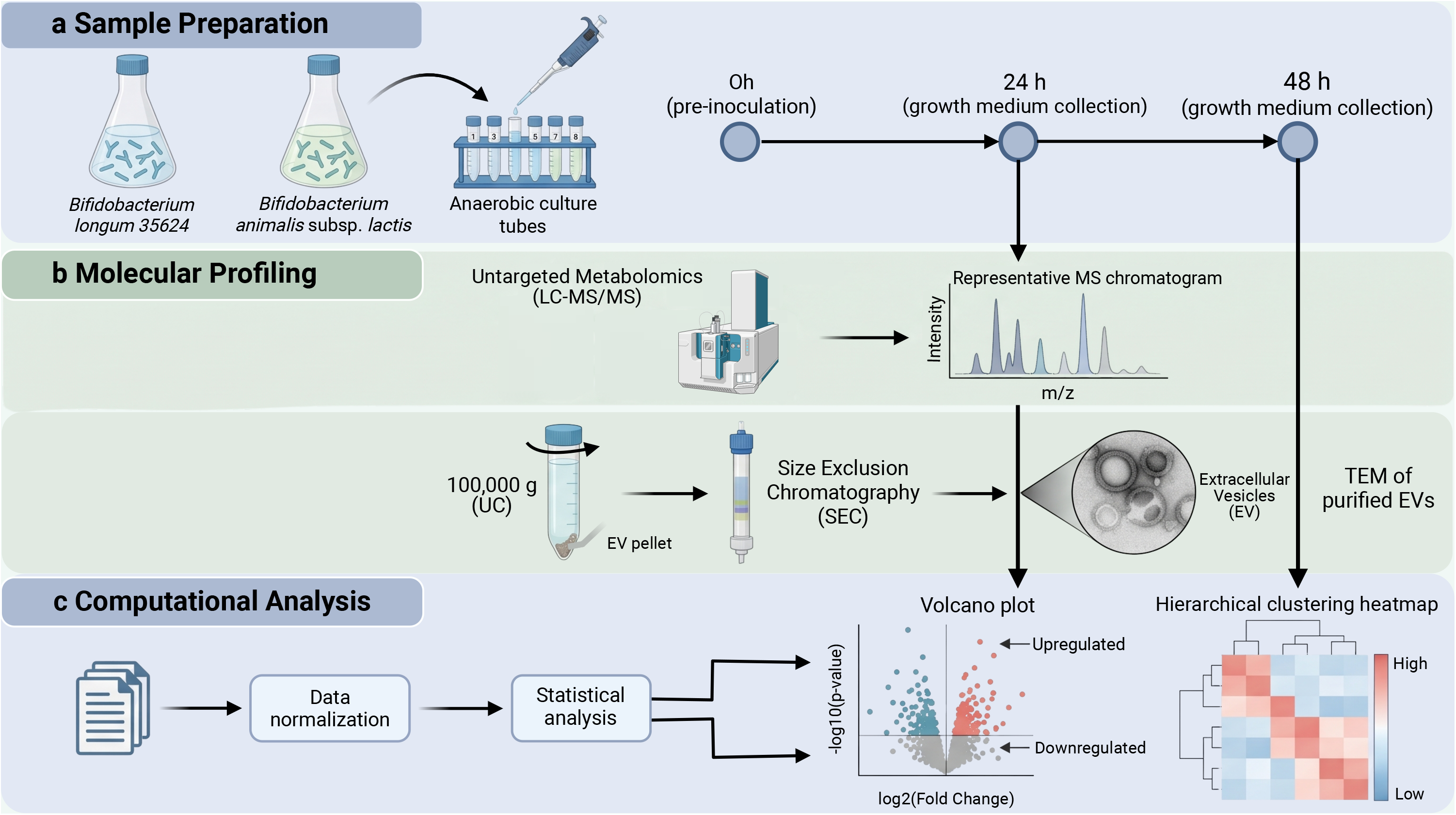
Study design and bacterial culture workflow. Schematic overview of the experimental design, anaerobic cultivation of the two *Bifidobacterium* strains, sample collection, vesicle isolation by differential ultracentrifugation, purification by size-exclusion chromatography and downstream untargeted LC–MS/MS metabolomics.

EVs were isolated from clarified supernatants by sequential centrifugation followed by differential ultracentrifugation, then purified by size-exclusion chromatography (SEC) to reduce contamination from soluble proteins and other extracellular macromolecules. Purified EVs from *B. animalis* subsp. *lactis* and *B. longum* 35624 were characterised by nanoparticle tracking analysis (NTA), time-resolved fluorescence immunoassay (TRFIA) and transmission electron microscopy (TEM) following MISEV2023 recommendations^14^ (Fig. 2). NTA gave particle concentrations of 1.2 × 10^11^ particles/mL for *B. animalis* subsp. *lactis* EVs and 5.6 × 10^10^ particles/mL for *B. longum* 35624 EVs in the final preparations (Fig. 2a). The two preparations had similar size distributions, with mode diameters of 107 nm (*B. animalis* subsp. *lactis*) and 109 nm (*B. longum* 35624), and most particles were between 50 and 300 nm (Fig. 2a). To define the marker profile of the vesicles, the bacterial EVs and EVs from the human prostate cancer cell line DU145 were analysed in parallel with two sandwich TRFIAs: LTA–LTA for the Gram-positive bacterial marker lipoteichoic acid (LTA) and CD63–CD63 for the human EV marker CD63 (Fig. 2b). The two assays gave opposite patterns. In the LTA–LTA assay, EVs from *B. animalis* subsp. *lactis* and *B. longum* 35624 gave high signals (signal-to-background ratio, S/B, 13.5 and 20.1), whereas DU145 EVs remained close to background (S/B 2.0), as expected for an assay that targets a bacterial cell envelope component. In the CD63–CD63 assay, DU145 EVs gave a high signal (S/B 58.4), whereas EVs from *B. animalis* subsp. *lactis* and *B. longum* 35624 remained close to background (S/B 2.0 and 1.6), as expected for a human tetraspanin that bacteria do not express. Each assay therefore detected its target only in the expected sample type. The bacterial EVs carried LTA without detectable CD63, consistent with LTA as a marker of Gram-positive bacterial membrane vesicles^15,16^, and the assay pair discriminated bacterial from human EVs.

**Fig. 2.**
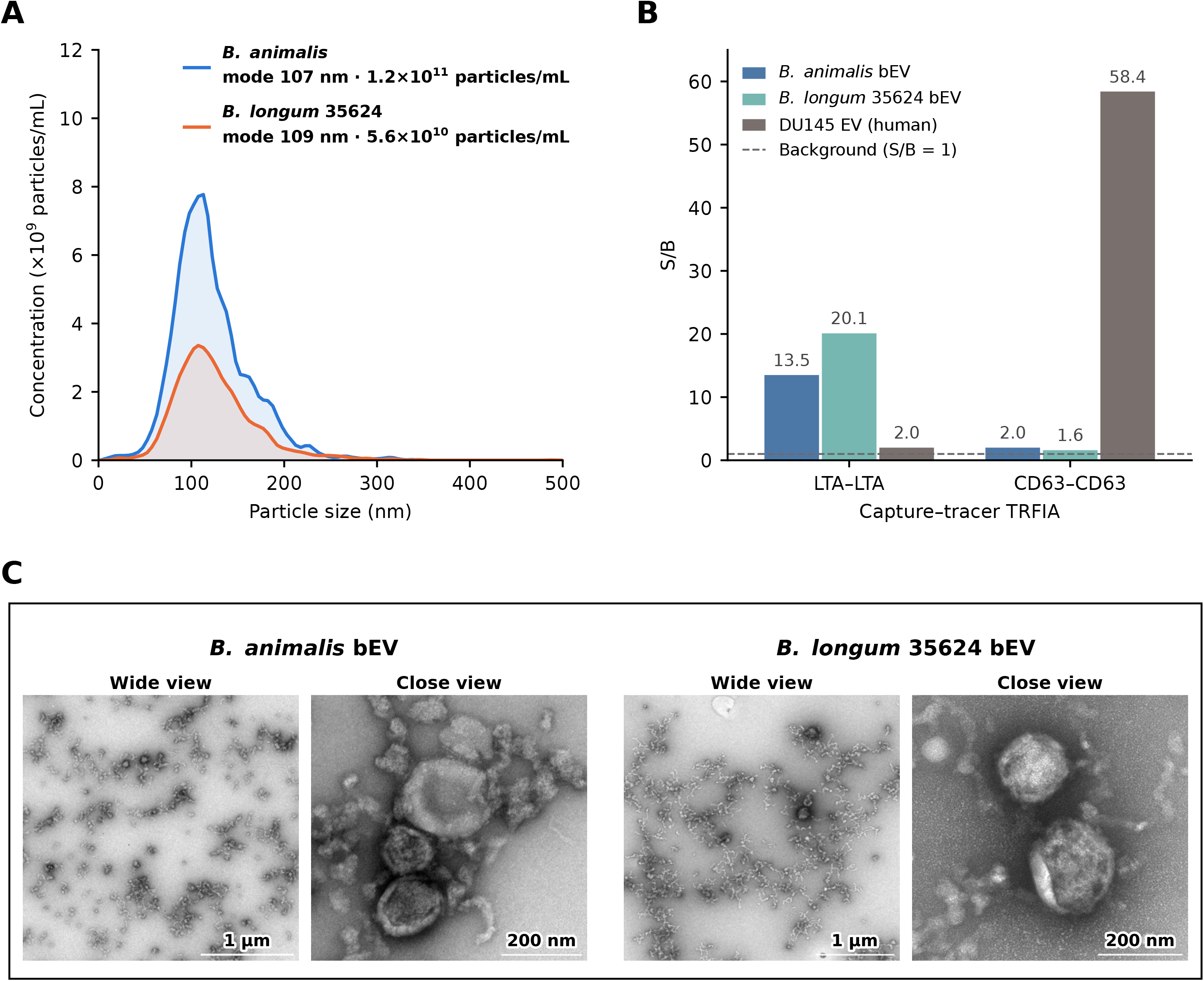
Characterisation of *Bifidobacterium*-derived extracellular vesicles (EVs). **a**, Size distribution of EVs from *B. animalis* subsp. *lactis* and *B. longum* 35624 measured by nanoparticle tracking analysis (NTA). **b**, Sandwich time-resolved fluorescence immunoassays (TRFIAs) for the Gram-positive bacterial marker lipoteichoic acid (LTA) and the human EV marker CD63, with homologous capture–tracer antibody pairs (1 µg total protein per well). DU145 cell-derived EVs served as the human EV reference (CD63-positive, LTA-negative). **c**, Representative negative-stain transmission electron microscopy (TEM) images of EVs from *B. animalis* subsp. *lactis* and *B. longum* 35624, each shown as a wide view and a close view (close-view scale bars, 200 nm). S/B, signal-to-background ratio.

The morphology of the purified EVs was examined by TEM. Images of EVs from both strains showed round, membrane-delimited vesicles consistent with the morphology of bacterial EVs (Fig. 2c). Together, the NTA, TRFIA and TEM results indicate that EVs were successfully isolated from both strains.

### Untargeted metabolomics: vesicles carry a sparse small-molecule cargo

Untargeted LC–MS/MS profiles of culture supernatants, EVs, uninoculated medium and solvent blanks were processed with the Everything Bagel feature finder 17 and analysed by feature-based molecular networking^18^ on GNPS2^19^ (Fig. 3a). Node pie charts in the network show the relative abundance of each feature across the EVs and the 24 h and 48 h supernatants of both strains and the medium. The enlarged molecular families illustrate structurally related metabolites that differed between sample groups, some being dominated by one sample group and their network neighbours by another (Fig. 3a). The supernatants of both strains contained a chemically diverse repertoire of amino acid, peptide, bile acid-related, fatty acid and amine conjugates; di- and tripeptide conjugates and multiplexed-synthesis conjugate library entries were the largest annotated classes (Supplementary Fig. S1), together with many molecular families that remain unannotated. Vesicles, by contrast, carried a sparse cargo. To assess the magnitude of the analytical background before any filtering was applied, we first performed principal component analysis on the unfiltered feature table: the EV samples of both strains co-localised with the solvent blanks rather than with the medium and supernatants (first component, 47.7% of the variance; Fig. 3b).

**Fig. 3.**
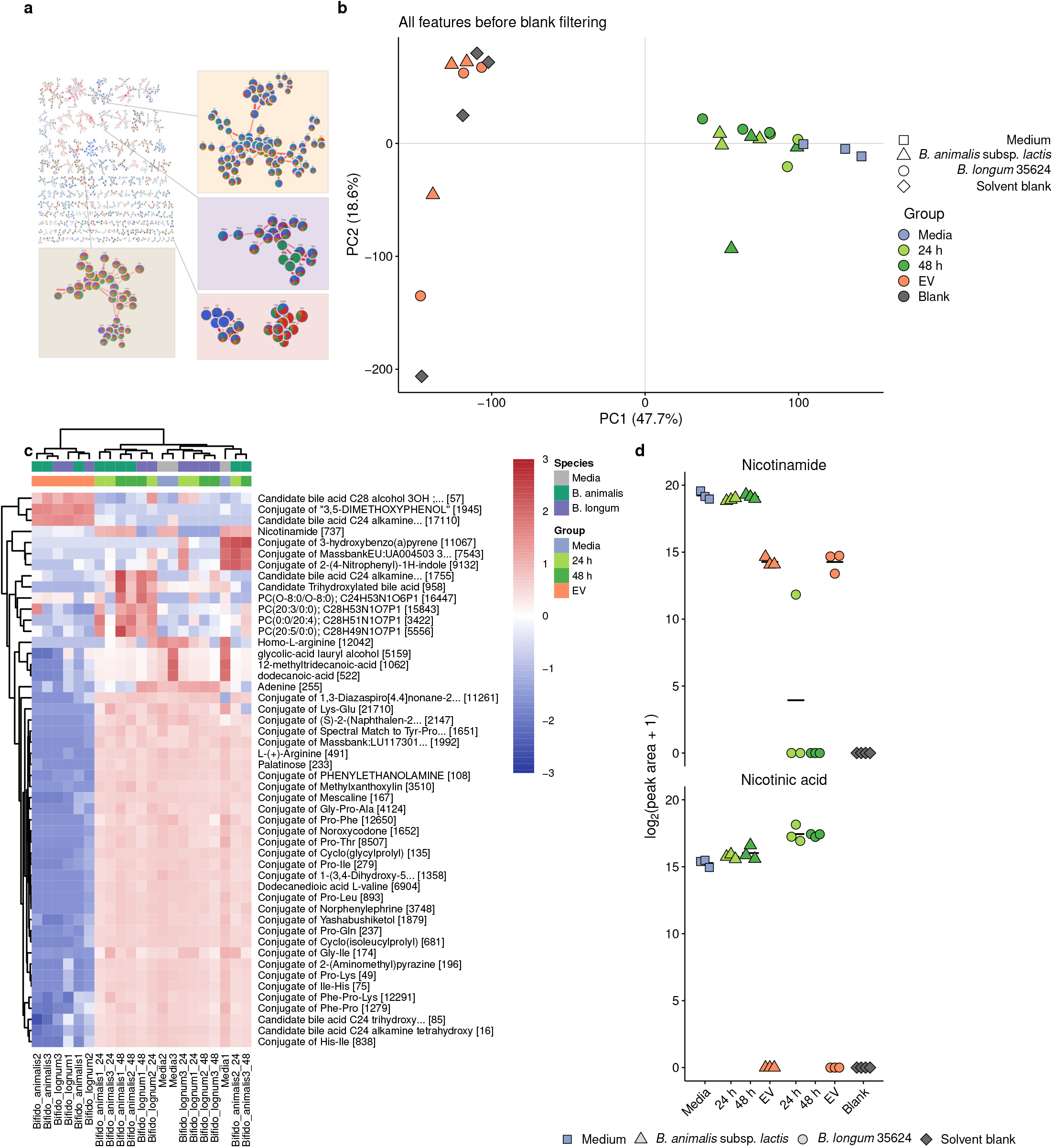
Untargeted metabolomes of *Bifidobacterium* supernatants and vesicles: vesicle profiles resemble analytical blanks and nicotinamide is converted in a strain-specific manner. **a**, Feature-based molecular network (GNPS2) of MS/MS features from culture supernatants, extracellular vesicles (EVs) and medium controls processed with Everything Bagel; nodes represent MS/MS features and edges spectral similarity. Node pie charts show the relative abundance of each feature across six sample groups: *B. animalis* subsp. *lactis* EVs (blue), *B. animalis* subsp. *lactis* 24 h (red), *B. animalis* subsp. *lactis* 48 h (green), *B. longum* 24 h (orange), *B. longum* 48 h (purple) and *B. longum* EVs (teal); the uninoculated medium was not included in the pie charts. Enlarged insets show selected molecular families in which structurally related features are most abundant in different sample groups. b, Principal component analysis of all features before blank filtering, including the solvent blanks (*n* = 4); zero values were replaced by half the feature minimum and data were log_2_-transformed and autoscaled. The EV samples of both strains co-localise with the blanks rather than with the medium and culture supernatants, indicating that vesicles carry few metabolites above background. c, Hierarchical clustering of the 50 most variable annotated features after blank filtering (row values: autoscaled log_2_ abundance; complete linkage, Euclidean distance, applied to features and samples). EVs separate from the medium and supernatants mainly through the absence of medium-derived metabolites, with a small set of features higher in EVs. Column bars show condition and species; numbers in square brackets are GNPS query scans. d, Nicotinamide and nicotinic acid (log₂(peak area + 1)) in the uninoculated medium, the 24 h and 48 h supernatants and the EVs of each strain, and in solvent blanks; nicotinamide is depleted and nicotinic acid rises only in *B. longum* 35624. Triangles, *B. animalis* subsp. *lactis*; circles, *B. longum*; squares, medium; diamonds, solvent blanks; horizontal bars show the group mean. The MS/MS library match for nicotinamide is shown in Supplementary Fig. S3d. n = 3 biological replicates per group (n = 4 solvent blanks).

Blank filtering was performed using the FBMN-STATS criterion^20^ and additionally excluded polyethylene glycol (PEG) annotations (Supplementary Fig. S2). After filtering, the EV samples of both strains clustered together and were separated from the intermixed medium and supernatant samples mainly by the absence of medium-derived metabolites (Fig. 3c). To characterise the vesicle metabolome, the EVs of each strain were compared with the medium and with the 24 h and 48 h supernatants, and features were ranked by the consistency of their difference across the three comparisons. Candidate bile acid conjugates were prominent among the EV-enriched features (Fig. 4a, b): seven of the 25 top-ranked features in *B. animalis* subsp. *lactis* and five in *B. longum* were candidate bile acids, four of them shared by both strains, including amine-conjugated (alkamine) and hydroxylated C24 bile acids and, in *B. animalis* subsp. *lactis*, a trihydroxylated C28 bile alcohol. These assignments are based on MS/MS spectral similarity to library candidates (Supplementary Fig. S3a) and were graded according to Metabolomics Standards Initiative guidelines^21^ against propagated bile acid libraries^22^. Fatty acid-derived conjugates including palmitic acid, *trans*-13-octadecenoic acid and monolaurin conjugates and lipid species such as SM(d16:1/16:0) were also among the EV-enriched features. A further set of EV-enriched features was annotated as conjugates of industrial compounds such as caprolactam and 3,4-dimethylbenzoic acid, which were detected only in vesicles (Fig. 4c, d).

**Fig. 4.**
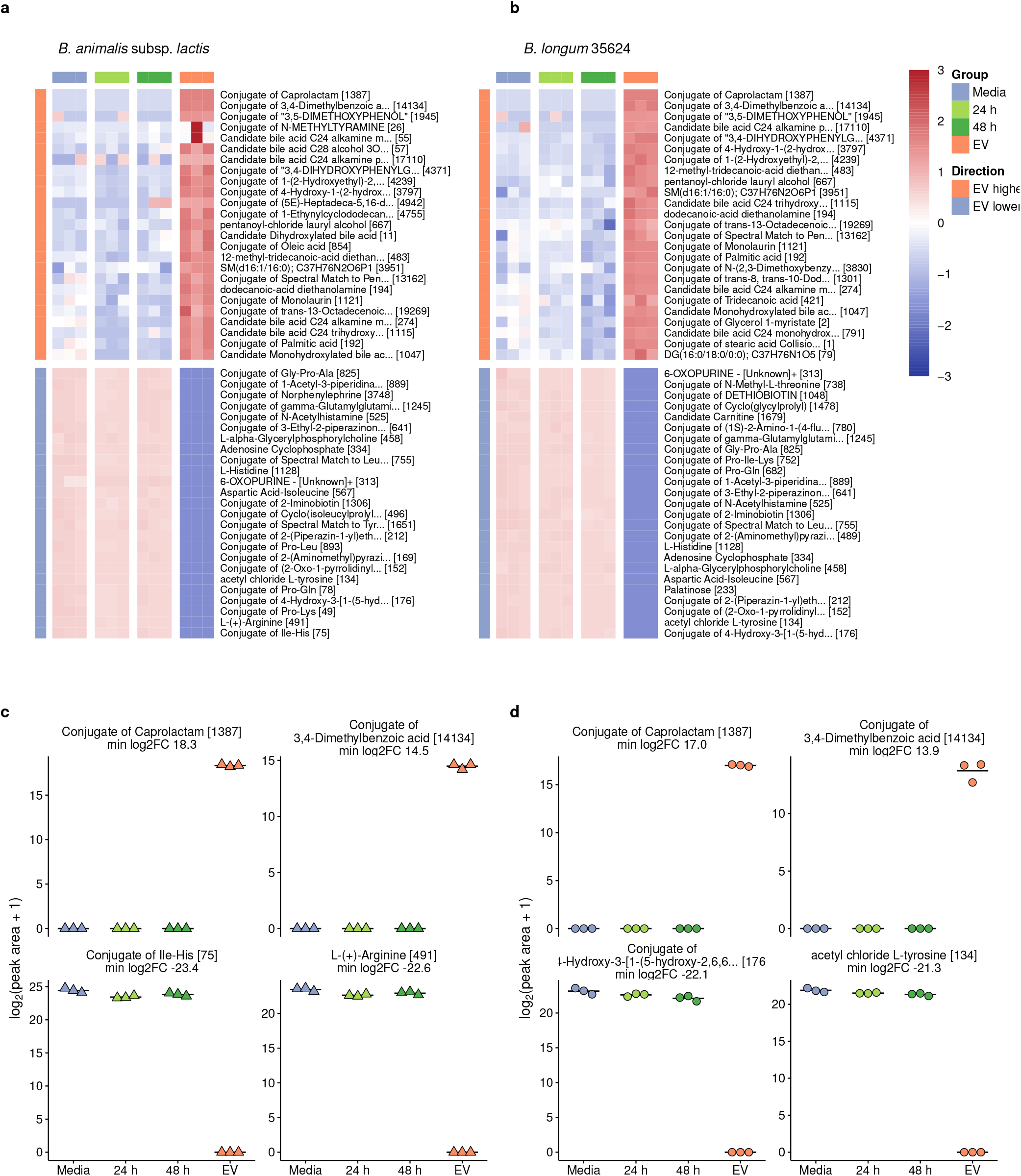
Metabolites associated with extracellular vesicles of *B. animalis* subsp. *lactis* and *B. longum*. **a,b**, The 25 annotated features most consistently higher (top) and the 25 most consistently lower (bottom) in EVs than in the uninoculated medium and in 24 h and 48 h culture supernatants of *B. animalis* subsp. *lactis* (**a**) and *B. longum* 35624 (**b**). Features were ranked by the smallest of the three log_2_ fold changes; values are row *z*-scores of log_2_(peak area + 1) across the strain’s samples, truncated at ±3. The side bar marks features higher or lower in EVs than in all three references; no comparison reaches FDR < 0.05 with n = 3. **c**,**d**, Individual log_2_(peak area + 1) values for the two most EV-enriched and the two most EV-depleted features of *B. animalis* subsp. *lactis* (**c**) and *B. longum* (**d**); horizontal bars show the group mean. Panel titles give the smallest of the three log_2_ fold changes; very large values indicate features not detected in the comparator groups. Numbers in square brackets are GNPS query scans. *n* = 3 biological replicates per group; comparisons are exploratory and test statistics are given in Supplementary Table 3.

The features most depleted from vesicles were medium nutrients and their derivatives (Fig. 4a, b): free amino acids (L-arginine and L-histidine), di- and tripeptides (for example Pro-Gln, Pro-Leu, Ile-His and Gly-Pro-Ala), cyclic dipeptides, acetylated amines and amino acids, glycerophosphocholine, cyclic AMP and hypoxanthine. These were abundant in the medium and in both supernatants but absent from, or near the detection limit in, vesicles (Fig. 4c, d). Such amino acids and amines are the building blocks of *N*-acyl amides and amine-conjugated bile acids^23–26^, a pattern consistent with conjugation products, rather than their precursors, being associated with vesicles. Across all 285 annotated features that passed the blank filtering and PEG filters, most were lower in vesicles than in the medium, and 31 per strain were higher (Supplementary Fig. S4). Our results also showed that the single vesicle-enriched feature with clear evidence of bacterial origin was a putative monohexosyl diacylglycerol (28:0), bifidobacterial membrane lipids which was higher in vesicles than in the 48 h supernatants, by approximately 2-fold in B. animalis subsp. lactis and 13- to 24-fold in B. longum 35624 (Supplementary Fig. S3b, c), consistent with vesicles being membrane-derived. We also found that many of the remaining vesicle-enriched features corresponded to polymer additives, plasticisers and disinfectant-related amines introduced during isolation rather than to bacterial products (Supplementary Fig. S5), reinforcing the need for the blank controls applied throughout.

### Strain-specific remodelling of the soluble exometabolome

The untargeted data also resolved how each strain reshaped the soluble medium over 48 h. Neither strain broadly depleted amino acids, di- and tripeptides or other nitrogen-rich nutrients; both grew largely on carbohydrate in this rich broth and were distinguished instead by a small number of specific conversions (Supplementary Fig. S6; Methods). Test statistics for these comparisons are given in Supplementary Table 2.

The clearest strain-specific conversion involved vitamin B3. In *B. longum* 35624, nicotinamide fell below the detection limit in two of three cultures by 24 h and in all cultures by 48 h, whereas nicotinic acid rose approximately 4-fold; in *B. animalis* subsp. *lactis*, nicotinamide was unchanged and nicotinic acid was only marginally higher (Fig. 3d; MS/MS library match in Supplementary Fig. S3d). This precursor–product pattern corresponds to deamidation of the nicotinamide amide group and is consistent with a nicotinamidase (PncA-type) activity expressed by *B. longum* 35624 but not detectably by *B. animalis* subsp. *lactis* in this medium. Both vesicle preparations contained only low levels of nicotinamide (about 3% of the medium level) and no nicotinic acid. The conversion was complete and strain-specific and provides a candidate functional marker of strain 35624 in culture. A second *B. longum*-specific product was a feature whose putative annotation suggests indole-3-lactic acid, a tryptophan catabolite characteristic of breast-milk-associated bifidobacteria^40^. This feature was absent from the medium, vesicles and solvent blanks, was detected only at trace levels in *B. animalis* subsp. *lactis*, and approximately doubled from 24 to 48 h in *B. longum* (Supplementary Fig. S7a). Free tryptophan remained abundant in all supernatants, consistent with conversion of only a small fraction of the available pool. The MS/MS spectrum of this feature contained the indole side-chain and water-loss fragments expected for indole-3-lactic acid but also co-eluting signals, so the annotation (MSI level 2–3) requires confirmation with an authentic standard and a negative-mode run to separate the phenyllactic and 4-hydroxyphenyllactic analogues.

The strains also handled purines in opposite directions. *B. animalis* subsp. *lactis* depleted adenosine, guanosine and adenine (approximately 10- to 27-fold lower than in the medium at 48 h) and accumulated hypoxanthine, whereas *B. longum* changed the nucleosides comparatively little, raised adenine and lowered hypoxanthine (Supplementary Fig. S7b). The adenosine family was also among the most strongly strain-differentiated clusters in the molecular network (Fig. 3a). The two strains would therefore present different purine pools to the host and to neighbouring microbes, although the responsible enzymes were not identified and one medium replicate was high for hypoxanthine. Finally, the strains used different carbohydrates from the same broth. A putative trisaccharide (the annotation suggests a hexose trimer) was cleared, and a putative free hexose strongly depleted, by *B. longum* within 24 h, whereas *B. animalis* subsp. *lactis* left the hexose and slightly accumulated the trisaccharide (Supplementary Fig. S7c); the dominant putative disaccharide pool was unchanged in both. These sugar annotations rest on accurate mass and adduct pattern and remain putative pending authentic standards. Together, these conversions a vitamin B3 deamidation and an aromatic lactate in *B. longum*, and nucleoside and adenine catabolism in *B. animalis* subsp. *lactis* — define a compact set of strain-specific exometabolome functions on a shared medium (Supplementary Fig. S8).

### Semi-targeted analysis of conjugated *N*-acyl lipids

We next examined the conjugated metabolites in more detail. Using an in-house library with retention-time matching to authentic standards, we detected thirteen conjugated *N*-acyl lipids, in which a fatty acid is linked to an amine-containing group through an amide bond (Fig. 5). Blank filtering was applied first, to the raw normalised peak areas; only Ala-C16:0 was detected in the blank, at levels far exceeding those in the samples, and it was removed. The twelve remaining lipids were conjugates of arginine (C4:0–C12:0), lysine (C10:0 and C12:0), alanine (C12:0) and amylamine (C11:0, C15:0 and C17:0). Two distribution patterns were observed among the conjugated *N*-acyl lipids (Fig. 5a–d). The arginine and lysine conjugates were already present in the uninoculated medium, remained at similar levels in the 24 h and 48 h supernatants and were absent from, or very low in, the vesicles of both strains, indicating that they derive at least in part from the medium. By contrast, Ala-C12:0 and the amylamine conjugates were absent from the medium and associated with vesicles. Ala-C12:0 was detected only in vesicles, in all three *B. longum* and one of three *B. animalis* subsp. *lactis* replicates; amylamine-C17:0 was restricted to vesicles in *B. animalis* subsp. *lactis* but was also present, at higher levels, in the 24 h *B. longum* supernatant; and amylamine-C11:0 was found at very low abundance in the vesicles of both strains and sporadically in 48 h supernatants. As for the untargeted data, these comparisons are exploratory (Supplementary Table 3).

**Fig. 5.**
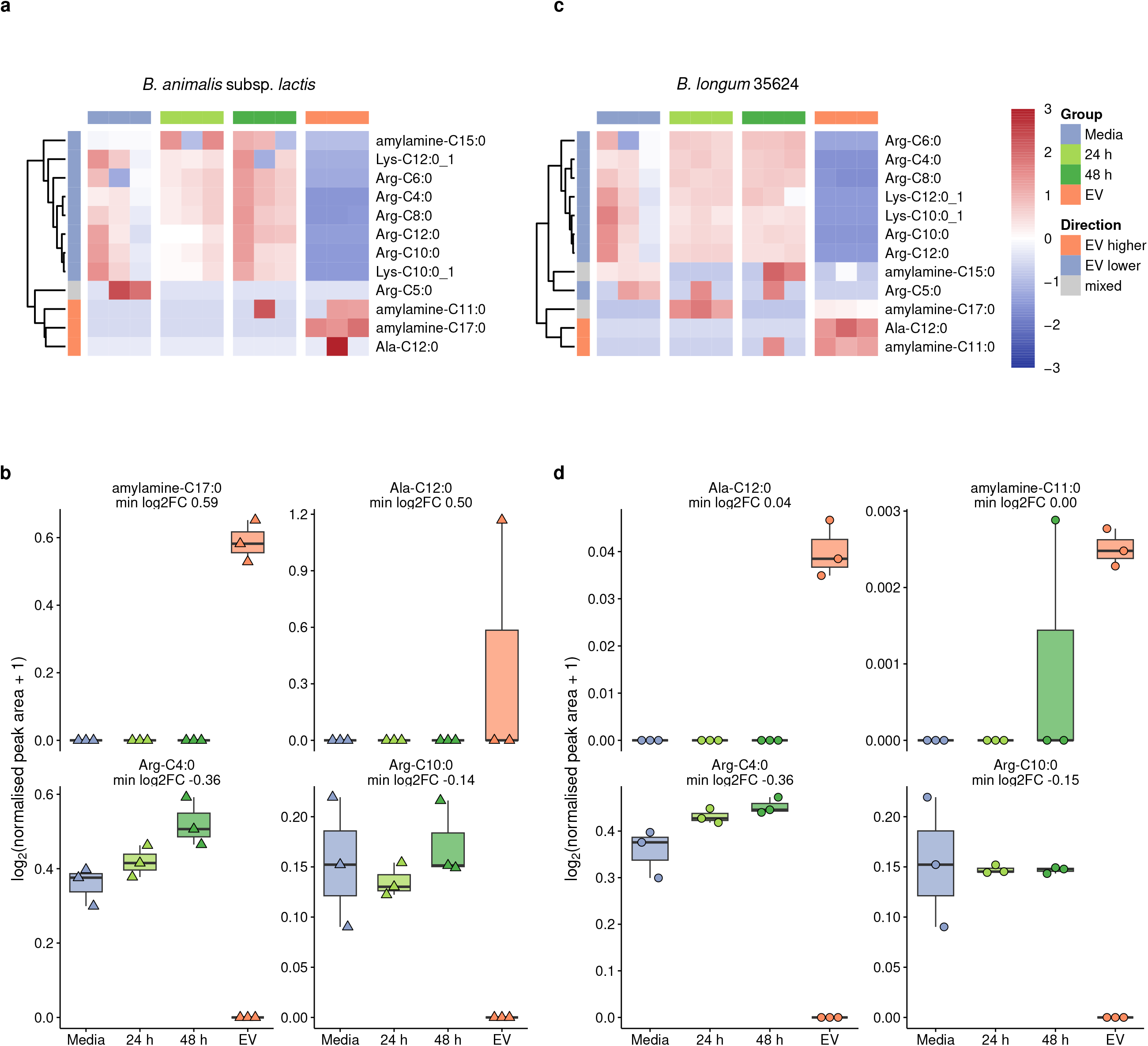
Conjugated *N*-acyl lipids in the *Bifidobacterium* exometabolome and EV metabolome (semi-targeted analysis). Thirteen conjugated *N*-acyl lipids were detected; one (Ala-C16:0) was present in the blank and removed, and the remaining twelve are shown. **a**,**c**, Row *z*-scores of log_2_(normalised peak area + 1) across the medium, 24 h and 48 h supernatants and EVs of *B. animalis* subsp. *lactis* (**a**) and *B. longum* 35624 (**c**); rows are clustered by complete linkage on Euclidean distances. The side bar classifies each lipid as higher in EVs than in all three comparators, lower than in all three, or mixed. **b**,**d**, Normalised peak areas (log_2_(*x* + 1)) of the two most EV-enriched and the two most EV-depleted lipids for *B. animalis* subsp. *lactis* (**b**) and *B. longum* (**d**). Box plots show the median (centre line), first and third quartiles (box bounds) and 1.5× the interquartile range (whiskers), with individual data points overlaid. Panel titles give the smallest of the three log_2_ fold changes, which are attenuated for these low-abundance lipids by the pseudocount of 1. *n* = 3 biological replicates per group; comparisons are exploratory and test statistics are given in Supplementary Table 3

## Discussion

This study provides a description of the exometabolome and the extracellular vesicle metabolome of two probiotic *Bifidobacterium* strains. The culture supernatants contained a chemically diverse repertoire of metabolites, including numerous conjugates, whereas vesicle profiles closely resembled analytical blanks. Most metabolites present in the medium were absent from vesicles, and only a small, selective set of candidate bile acid, fatty acid and *N*-acyl lipid conjugates was associated with them above background. To our knowledge, the small-molecule cargo of bifidobacterial vesicles, and in particular the emerging class of conjugated microbial metabolites, has not been reported before.

The soluble exometabolome revealed a compact set of strain-specific functions that extend bifidobacterial metabolism beyond carbohydrate fermentation (Fig. 3d and Supplementary Fig. S7). *B. longum* 35624 deamidated nicotinamide to nicotinic acid, a reaction attributable to a bacterial nicotinamidase; because host tissues can use microbially generated nicotinic acid through the deamidated NAD salvage route^41,42^, such a strain-specific step could in principle influence host NAD precursor supply. The same strain produced indole-3-lactic acid, an aromatic amino acid catabolite of breast-milk-associated bifidobacteria that engages the aryl hydrocarbon receptor and HCA3 in immune cells^40^, linking strain 35624 to an established host-facing signal. *B. animalis* subsp. *lactis* instead catabolised purine nucleosides and adenine while exporting hypoxanthine, and the two strains cleared different carbohydrates, suggesting complementary, potentially cross-feeding niches on a shared substrate. These conversions are individually simple, strain-specific and inexpensive to confirm with authentic standards, and we regard them as the most robust biological signal in the dataset.

Bacterial EVs were isolated by ultracentrifugation and further purified by SEC to reduce soluble non-vesicular material. NTA and TEM showed particles with the size and morphology expected for EVs, and the two TRFIAs gave complementary marker profiles: the Gram-positive bacterial marker LTA was detected on EVs from both strains but not on human DU145 EVs, whereas the human EV marker CD63 was detected on DU145 EVs but not on the bacterial EVs. These findings indicate that bacterial EVs were present in the analysed preparations, so the blank-like vesicle metabolite profiles are unlikely to reflect failed isolation. The similarity between vesicles and blanks instead suggests that these EVs carry a sparse cargo within a very small luminal volume, which is biologically plausible. Candidate bile acid conjugates, including amine-conjugated (alkamine) C24 bile acids shared by both strains, were the most consistent vesicle-enriched features however, could not be attributed to bacterial production, as they were present in the culture medium. Microbially conjugated bile acids engage host bile acid receptors^3^, can influence immune cell differentiation and have been associated with islet autoimmunity^27^; moreover, bile salt hydrolases, which are widespread among bifidobacteria, also act as acyltransferases that conjugate bile acids to amino acids and amines^25,26^, providing a plausible route to the candidates observed here. Because bifidobacteria cannot synthesise the sterol nucleus, such conjugates would require bile acid or sterol substrates from the complex medium, in which other bile acid-like features were indeed detected. These annotations are tentative, and their structures, origin and vesicle association require confirmation with authentic standards and isotope tracing.

Conjugated *N*-acyl lipids are a dynamic component of microbial chemistry. We recently observed that microbiome diversifies the *N*-acyl lipid pool across a wide range of acyl chain lengths^24^, and microbially conjugated lipids shift systematically between compartments of a simulated human gut^28^, yet in neither setting can a conjugate be attributed to a single producer. Profiling axenic cultures alongside sterile medium allowed this distinction and proved essential. The arginine and lysine conjugates, which formed the majority of the lipids detected, were already present in the uninoculated medium and cannot be attributed to bifidobacterial synthesis, whereas Ala-C12:0 and the amylamine conjugates were absent from the medium and associated with vesicles, making them the strongest candidates for bacterially derived *N*-acyl lipids. Their distribution differed between strains: amylamine-C17:0 was restricted to vesicles in *B. animalis* subsp. *lactis* but appeared transiently in the 24 h *B. longum* supernatant, suggesting selective, strain-specific loading, although pooling of vesicles across the 24 h and 48 h harvests precludes conclusions about kinetics. Conversely, amino acids, peptides and amines the building blocks of *N*-acyl amides and amine-conjugated bile acids^23–26^ were abundant in the medium and supernatants but absent from vesicles. We therefore hypothesise that bifidobacteria draw on this extracellular pool to form conjugates, a subset of which becomes associated with vesicles; supplying stable-isotope-labelled substrates in a defined medium would test this directly.

A further limitation concerns the conjugated metabolites associated with vesicles. The cultures showed an overall, non-specific drop in signal intensity relative to the uninoculated medium; once this was taken into account (Methods), neither the candidate bile acid class nor the *N*-acyl amino acid class showed a coherent production pattern in either strain, and several conjugates including the arginine and lysine *N*-acyl lipids were in part already present in the uninoculated medium (Supplementary Fig. S6). These conjugates are therefore reported as candidates detected alongside vesicles rather than as confirmed bacterial products, and their origin remains to be established. A late rise in free cholic acid at 48 h was likewise too variable between replicates (approximately 100-fold) to interpret (Supplementary Fig. S6). The clearest bacterial signal in the vesicle fraction was consequently a membrane glycolipid rather than a conjugated small molecule, consistent with bifidobacterial vesicles being membrane-derived^43^ and with most of their lipid cargo lying beyond the mass range measured here.

Our findings do extend the view of bifidobacterial metabolism beyond classical fermentation products^5^. *B. longum* 35624 has well-documented immunomodulatory properties^29^, *B. animalis* subsp. *lactis* interacts with the host epithelium, immune system and resident microbiota and supports barrier integrity^30,31^, and bacterial vesicles can modulate host barrier and immune function in vivo^8–10,44^. If selective packaging of conjugates is confirmed, vesicles would allow bifidobacteria to deliver defined, potentially receptor-active metabolites^23^ to host or microbial targets in a protected form a hypothesis generated by descriptive data that remains to be tested. This exploratory study has several limitations. Blank filtering was essential to prevent background signals, including PEG-related polymers and candidate bile acids and lipids, from being misidentified as vesicle cargo. Although this approach may also have excluded genuine vesicle-associated metabolites, it highlights the importance of process blanks, transparent filtering criteria and the reporting of excluded features in EV metabolomics. Our study was limited to two strains grown in a single rich medium, with a small sample size that limits statistical power; the findings should therefore be considered exploratory. Although NTA, TRFIA and TEM supported EV identification after ultracentrifugation and SEC, co-isolated non-vesicular material was not quantified, and NTA does not distinguish vesicles from similarly sized non-vesicular particles. Some of the metabolite annotations remain putative (MSI level 2) or tentative (MSI level 3) and require confirmation with reference standards, absolute quantification and functional validation. Several strongly vesicle-enriched features were annotated as conjugates of industrial compounds, including caprolactam, 3,4-dimethylbenzoic acid and fatty acid diethanolamide surfactants; their apparent vesicle specificity may reflect contamination introduced during isolation, which solvent blanks alone cannot exclude. Finally, the limited vesicle material and the lack of normalisation to particle number or protein content may have reduced metabolite detection and preclude quantitative comparisons of abundance per vesicle. In summary, two probiotic *Bifidobacterium* strains release a chemically diverse exometabolome, whereas their vesicles carry a sparse cargo that closely resembles analytical blanks and is distinguished mainly by the absence of medium-derived metabolites and by a small, selective set of candidate bile acid, fatty acid and *N*-acyl lipid conjugates. Rigorous blank control is essential for interpreting such low-biomass samples; however, our study is the first map that provides a future reference for mechanistic and in vivo studies.

## Methods

### Bacterial strains and anaerobic culture

*Bifidobacterium longum* 35624 (PrecisionBiotics, UK) and *Bifidobacterium animalis* subsp. *lactis* (Winclove Probiotics, The Netherlands) were cultivated under strictly anaerobic conditions in a Whitley A35 Anaerobic Workstation (Don Whitley Scientific, UK). A 10 µL aliquot of bacterial glycerol stock stored at −80 °C was aseptically transferred into 2 mL Fastidious Anaerobe Broth (FAB; Neogen, USA). Three independent biological replicates were prepared per strain. Samples were collected from uninoculated medium (medium control), immediately after inoculation (0 h), and after 24 h and 48 h of incubation. Growth was monitored by optical density at 600 nm. Cell-free supernatants were obtained by centrifugation and stored at −80 °C until metabolomic analysis. For extracellular vesicle isolation, parallel cultures were prepared by inoculating 100 µL of bacterial stock into 50 mL FAB. Three biological replicates were established per strain together with uninoculated medium controls. Vesicle-containing supernatants were collected after 24 h and 48 h of incubation and pooled across the two harvest times to provide sufficient material for characterisation and metabolomic analysis.

### Whole-genome sequencing

Whole-genome sequencing was performed to confirm strain identity. Genomic DNA was extracted from pure cultures (NucleoSpin Microbial DNA Mini kit, Macherey-Nagel, Germany). Sequencing libraries were prepared with the Nextera XT kit (Illumina) according to the manufacturer’s instructions and sequenced on an Illumina MiniSeq platform using 150 bp paired-end reads. Raw reads underwent adapter removal and quality filtering, followed by species confirmation, de novo assembly and annotation using CLC Genomics Workbench 26.0.1 (Find Best Match using K-mer Spectra, De Novo Assemble Small Genome, Find Prokaryotic Genes, and Annotate CDS with Best DIAMOND Hit). The best k-mer match for the *B. animalis* isolate was *Bifidobacterium animalis* subsp. *lactis*. Software versions and parameters, together with assembly statistics, are given in Supplementary Table 4.

### Isolation of bacterial extracellular vesicles

Vesicles were isolated by differential centrifugation and ultracentrifugation with minor modifications of a published protocol^32^. Cells were pelleted twice at 4,500 × *g* for 20 min at 4 °C to remove intact cells and large debris, followed by centrifugation at 10,000 × *g* for 30 min at 4 °C (Avanti J-26S XP with JA-14 fixed-angle rotor; Beckman Coulter). Supernatants were filtered through 0.22 µm low-protein-binding membranes (Filtropur S, Sarstedt, Germany). Clarified supernatants were ultracentrifuged at 150,000 × *g* for 2 h at 4 °C (Type 45 Ti rotor, Optima XE-90; Beckman Coulter); pellets were washed with 0.22 µm-filtered cold phosphate-buffered saline (PBS) and ultracentrifuged again under the same conditions. Enriched vesicle pellets were resuspended in PBS and stored at −80 °C. A PBS process blank was carried through the complete workflow in parallel.

### Size-exclusion chromatography

To improve vesicle purity and minimise contamination from soluble proteins and extracellular macromolecules, ultracentrifugation-derived preparations were further purified by size-exclusion chromatography using qEV columns (qEVoriginal 70 nm; Izon Science, Christchurch, New Zealand)^32^. Suspensions were applied to pre-equilibrated columns according to the manufacturer’s instructions and sequential fractions were collected. Vesicle-containing fractions were identified from their elution profile and pooled for downstream analysis^32^. Pooled fractions were concentrated where necessary and stored at −80 °C prior to electron microscopy and metabolomic analysis. Column manufacturer, model, pore size, buffer, loading volume and the identity of the pooled fractions are specified in the Supplementary Methods.

### Nanoparticle tracking analysis

The particle concentration and size distribution of EVs from *B. animalis* subsp. *lactis* and *B. longum* 35624 were measured by NTA as described previously^33,34^. A ZetaView PMX-120 instrument (Particle Metrix, Ammersee, Germany) with a 488 nm laser was used in scatter mode. The bacterial EV samples were diluted in PBS filtered through a 0.22 µm membrane before measurement (*B. animalis* subsp. *lactis* 1:2000, *B. longum* 35624 1:1000). Videos were recorded at 11 positions over 3 cycles, with sensitivity 85, shutter 100, a frame rate of 30 fps and a temperature of 22 °C. The data were analysed with ZetaView software version 8.06.01 SP1 using minimum brightness 30, minimum area 10, maximum area 1000, trace length 15 and a 5 nm size bin. Particle concentrations were corrected for the dilution factor, and the mode diameter and total particle concentration (particles/mL of EV preparation) were reported for each sample.

### Time-resolved fluorescence immunoassay

Two sandwich TRFIAs with homologous capture–tracer antibody pairs were used to define the marker profile of the EVs. Lipoteichoic acid (LTA), a cell envelope glycolipid of Gram-positive bacteria that is carried by their membrane vesicles^15,16^, served as the positive marker for EVs from *B. animalis* subsp. *lactis* and *B. longum* 35624, whereas the human tetraspanin CD63 served as the negative marker for the same EVs. EVs from the DU145 prostate cancer cell line were used as the human EV reference, providing the positive sample for the CD63–CD63 assay and the negative sample for the LTA–LTA assay, so that each assay had its own positive and negative samples. The mouse monoclonal anti-LTA antibody (IgG1, clone G43J; cat. no. MA1-7402) was purchased from Invitrogen (Thermo Fisher Scientific, Waltham, MA, USA). The mouse monoclonal anti-CD63 antibody (IgG1κ, clone H5C6; cat. no. 556019) was purchased from BD Pharmingen (BD Biosciences, Vantaa, Finland). Both assays followed our previously described nanoparticle-based protocol^34,35^. Biotinylated anti-LTA or anti-CD63 capture antibodies (100 ng/well) were diluted in red assay buffer (Uniogen Oy, Turku, Finland) and added to streptavidin-coated 96-well plates (KaiSA96, Uniogen Oy) at 30 µL/well. After 1 h incubation at room temperature (RT), the plates were washed twice with wash buffer (Uniogen Oy). The EV samples (1 µg total protein per well) were diluted in red assay buffer, added at 30 µL/well and incubated for 1 h at RT with slow shaking. After two washes, the tracer was added: Eu^3+^-nanoparticles coated with anti-LTA or anti-CD63 antibodies (1 × 10^7^ nanoparticles in 30 µL/well). After 1.5 h incubation at RT, the plates were washed four times. Time-resolved fluorescence was measured (λ_ex_ 340 nm, λ_em_ 615 nm) with a Victor 1420 multilabel counter (PerkinElmer, Turku, Finland). Wells containing buffer only, without EVs, served as background. All samples were measured in duplicate wells and the results are shown as signal- to-background ratio (S/B, mean ± SD).

### Transmission electron microscopy

Negative-stain transmission electron microscopy was performed to assess vesicle morphology, size and structural integrity in accordance with MISEV2023 guidance^14^. Carbon-coated 200-mesh copper grids were rendered hydrophilic by plasma cleaning (GloQube Plus, Quorum Technologies, UK). Purified samples were fixed in 2% paraformaldehyde for 2 h at room temperature. A 5 µL aliquot of fixed sample was deposited onto grids and incubated for 5 min. Grids were rinsed with filtered water, negatively stained with 1% uranyl acetate for 1 min, blotted and air-dried for 5 min. Imaging used a JEM-1400 microscope (JEOL, Tokyo, Japan) at 80 kV with an SIS Quemesa CCD camera (Olympus, Tokyo, Japan). Representative images were acquired from multiple fields of view across independent grids.

### Untargeted LC–MS/MS metabolomics

Methanol, water and acetonitrile were CHROMASOLV™ LC-MS Ultra grade (Honeywell, Riedel-de Haën, Charlotte, NC, USA) and ammonium acetate was LC-MS LiChropur® grade (Merck, Germany). Untargeted profiling of culture supernatants, extracellular vesicles and medium controls was performed by ultra-high-performance liquid chromatography coupled to high-resolution tandem mass spectrometry. Metabolites were extracted using a monophasic solvent procedure based on the SIMPLIFY workflow with modifications^36^, providing coverage of polar, semipolar and nonpolar metabolites and lipids from a single extract. Samples were analysed in randomised order together with pooled quality-control samples and solvent blanks (*n* = 4). A SCIEX ZenoTOF 8600 mass spectrometer operating in information-dependent acquisition (IDA) mode with positive ionisation was combined with a Waters Acquity Premier UHPLC system (Waters Inc., Milford, CT, USA). The stationary phase was a Waters Premier HSS T3 column (1.8 µm, 100 mm × 2.1 mm) fitted with a pre-column of the same material and held at 45 °C. The aqueous mobile phase was 2 mM ammonium acetate in water and the organic phase was 2 mM ammonium acetate in methanol, at a flow rate of 400 µL/min. The gradient started at 100% aqueous phase, the organic content was increased to 72% over 5.5 min and then to 100% over a further 7.5 min, and was held for 3.5 min. The injection volume was 0.2 µL, and the injection needle was washed for 8 s with 0.1% formic acid in methanol:acetonitrile:water:isopropanol (3:3:3:1). The autosampler was held at 15 °C. Ion source gas 1 was 60 psi, gas 2 was 80 psi, the curtain gas was 50 psi, the CAD gas was 8, the declustering potential was 40 V, the drying gas temperature was 350 °C and the spray voltage was 5500 V. IDA criteria were a mass tolerance of 50 mDa, a maximum of 10 candidate ions, a threshold of 3000 counts/s and dynamic background subtraction. The collision energy used for fragmentation was 40 V in CID mode. The sample-amount normalisation basis for vesicle extracts and the quality-control acceptance criteria are given in the Supplementary Methods.

### Semi-targeted analysis of conjugated *N*-acyl lipids

Conjugated *N*-acyl lipids were profiled using an in-house spectral and retention-time library constructed from authentic standards. Candidate features were matched on accurate precursor mass, characteristic amide-bond fragmentation and retention time. Peak areas were integrated in SCIEX OS version 4.2.0 and normalised peak areas were exported for all samples.

### Data processing and molecular networking

Raw data were processed with the Everything Bagel Feature Finder (version 2.0.0+v2)^17^, which detects chromatographic features from their mass-to-charge ratio (*m/z*) and retention time, quantifies each feature by its integrated peak area and removes signals below an adaptive MS1 noise threshold (1,165.34). Features detected in a minimum number of samples were aligned, and the consensus features with their peak areas and MS/MS spectra were used for all downstream analyses. MS/MS data were analysed by feature-based molecular networking^18^ on GNPS2^19^. Spectral library matching used the GNPS public libraries, including propagated bile acid^22^ and reverse-metabolomics conjugate^37^ libraries, and for each feature the match with the highest cosine score was retained; the linking of library matches to features is described in the Supplementary Methods. Compound classes were assigned to the annotated features by rule from the library annotation names (class group and fine class; Supplementary Table 3). Annotation confidence was assigned according to Metabolomics Standards Initiative guidelines^21^: direct library matches as level 2, and annotations from propagated or multiplexed-synthesis libraries (prefixed ‘Candidate’ or ‘Conjugate of’) as level 3. In figures, compound names are shortened and followed by the GNPS query scan number in square brackets. The GNPS2 job is available at https://gnps2.org/status?task=573b8bf50df649999443757db4c675e6.

### Statistical analysis

All data processing and statistical analyses were performed in R 4.4.1^38^ using readxl, readr, dplyr, tidyr and stringr for data handling, ggplot2 and ggbeeswarm for plotting, pheatmap for heatmaps, patchwork for figure assembly and openxlsx for the supplementary tables. All statistical results, Figs. 3–5 (except the molecular network in Fig. 3a), Supplementary Figs. S1–S8 and Supplementary Tables 1–5 are produced by the scripts from the input data (Code availability). The untargeted and semi-targeted datasets were processed with separate scripts (Code availability) but analysed with the same statistical workflow. For each strain, EV samples were compared with the uninoculated culture medium and with the 24 h and 48 h culture supernatants (*n* = 3 biological replicates per group); the same medium samples served as the reference for both strains.

#### Untargeted data

Peak areas were used without further normalisation, and missing values were treated as not detected (zero). Features with zero intensity in all samples and blanks were removed. Background features were then removed following FBMN-STATS^20^: a feature was discarded when (mean intensity in the solvent blanks + 1)/(mean intensity across all non-blank samples + 1) was ≥ 0.3. Every removed feature was recorded (Supplementary Methods; Supplementary Table 5). Statistical analyses were restricted to annotated features, and candidate PEG annotations were excluded before any statistical analysis. The numbers of features remaining after each step are given in Supplementary Table 5.

#### Semi-targeted data

Values reported as not detected in SCIEX OS were set to zero. Blank filtering was the first processing step and was applied to the untransformed normalised peak areas: a lipid was removed when the ratio of its mean in the blank to its mean across all non-blank samples was ≥ 0.3 (the FBMN-STATS criterion, without an offset), and lipids not detected in the blank were retained. Lipids not detected in any sample were excluded, and for each strain only lipids detected in at least one sample of that strain or of the medium were tested.

#### Univariate analysis

For each strain, EV samples were compared with each reference group (medium, 24 h and 48 h) using two-sided Wilcoxon rank-sum tests (normal approximation with continuity correction), and *P* values were adjusted within each comparison by the Benjamini–Hochberg procedure^39^. With three replicates per group the smallest attainable *P* value is 0.08 (0.06 in the presence of ties), so no comparison could reach nominal significance. Because the replicate number is small and the study is exploratory, the comparisons are reported as a discovery ranking rather than as confirmatory tests; *P* values are tabulated in Supplementary Table 3 but were not used to select metabolites. Log_2_ fold changes were calculated from group mean peak areas (untargeted) or group mean normalised peak areas (semi-targeted) using a pseudocount of 1, as log_2_[(mean EV + 1)/(mean reference + 1)]. Metabolites with a positive log_2_ fold change in all three comparisons were classified as EV-enriched and those with a negative log_2_ fold change in all three as EV-depleted; within each class, metabolites were ranked by the smallest absolute log_2_ fold change of the three comparisons, that is, by their minimum difference from any reference group. Because most normalised peak areas in the semi-targeted data were below 1, the pseudocount attenuates the fold changes of low-abundance lipids, and the reported semi-targeted fold changes are therefore conservative. Where several untargeted features carried the same annotation, all were tested, but only the highest-ranked feature per compound is shown in figures. No result is presented as statistically confirmatory.

#### Multivariate analysis

For principal component analysis (PCA) and hierarchical clustering, zero values were replaced by half the minimum detected value of each feature, and the data were log_2_-transformed and autoscaled (mean-centred and divided by the standard deviation of each feature). To assess background signal, PCA including the solvent blanks was performed on all features before blank filtering (Fig. 3b). Hierarchical clustering of features and samples used complete linkage on Euclidean distances of the autoscaled data and was applied to the 50 most variable annotated features after blank filtering (Fig. 3c).

#### Visualization

In the per-strain heatmaps, values are row *z*-scores of log_2_(intensity + 1) across that strain’s samples (medium, 24 h, 48 h and EV), with the colour scale truncated at ±3 for display. The untargeted heatmaps show the 25 most EV-enriched and the 25 most EV-depleted features, ranked as above and ordered by fold change (Fig. 4a, b); the semi-targeted heatmaps show all twelve lipids, with rows clustered by complete linkage on Euclidean distances (Fig. 5a, c). For selected features, individual log_2_(intensity + 1) values are shown as dot plots with the group mean and standard deviation (untargeted; Fig. 4c, d) or as box plots with individual data points (semi-targeted; Fig. 5b, d).

#### Strain-specific exometabolome analysis

All MS/MS features of the Everything Bagel feature table that passed the blank filter (5,172 features) were compared between groups by Welch’s t tests on log^2^(peak area + 1), with Benjamini–Hochberg correction across all retained MS/MS features (12 comparisons: each strain at 24 h and 48 h versus the medium, 48 h versus 24 h, the two strains at each time point, and the EVs of each strain versus its 48 h supernatant and versus the medium; Supplementary Table 2). To compare consumption and production independently of an overall, non-specific difference in signal intensity between the cultures and the uninoculated medium (arising, for example, from dilution, ion suppression or injection differences), each medium and culture sample was divided by its global signal shift factor, 2 to the power of its median log^2^ ratio to the mean of the medium over the 4,648 features detected in all 15 medium and culture samples, and the comparisons were repeated on the corrected values (Supplementary Fig. S6; Supplementary Table 2b,c). Individual metabolites (nicotinamide, nicotinic acid, indole-3-lactic acid, tryptophan, purine bases and nucleosides, and sugar features) were confirmed by accurate mass (within 10 ppm of the expected *m/z*) and MS/MS, and bifidobacterial membrane lipids (diacylglycerols and mono- and dihexosyl diacylglycerols) were searched for in the vesicle samples by the accurate masses of their sodium and ammonium adducts. For the class-level overview, EVs of each strain were compared with the medium by Welch’s t tests on log^2^(peak area + 1) for the annotated features that passed the blank and PEG filters (Supplementary Figs. S1 and S4; Supplementary Table 3).

## Data availability

The untargeted LC–MS/MS data generated in this study have been deposited in the GNPS/MassIVE repository under accession [MassIVE MSV number to be added]. The GNPS2 feature-based molecular networking job (Everything Bagel) is available at https://gnps2.org/status?task=573b8bf50df649999443757db4c675e6. The processed input data used for all analyses (Everything Bagel feature table, GNPS2 library matches, sample metadata and the SCIEX OS export of the semi-targeted data) are provided with the analysis code, and the processed results are given in Supplementary Tables 1–5.

## Code availability

Analysis code is available at https://github.com/LamichhaneSantosh/Bifidobacteria_Metabolomics and archived on Zenodo under DOI [Zenodo DOI to be added]. The repository is a self-contained R project with the input data: 01_import_and_blank_filter.R links GNPS2 library matches to features (with precursor *m/z* verification), records the fate of every library match and applies the blank filter; 02_EV_annotated_compound_analysis.R performs PEG exclusion, the EV comparisons, PCA and hierarchical clustering; 03_exometabolome_strain_specific.R performs the strain-specific exometabolome comparisons and the global signal shift correction; 04_compound_class_overview.R assigns compound classes and compares EVs with the medium; 05_SciexOS_Nacyl_lipids_statistics.R and 06_SciexOS_Nacyl_lipids_plots.R analyse the semi-targeted N-acyl lipids; and 07_main_figures.R, 08_supplementary_figures.R and 09_supplementary_tables.R produce Figs. 3–5, Supplementary Figs. S1–S8 and Supplementary Tables 1–5. 00_run_all.R runs the complete analysis.

## Acknowledgements

The authors thank the staff of the Institute of Biomedicine, University of Turku, Turku Bioscience Centre and the Turku Clinical Microbiome Bank for technical assistance and access to research infrastructure, and the metabolomics and electron microscopy core facilities for their expertise and support. Mass spectrometric analysis was performed at the Turku Metabolomics Centre with the support of Biocenter Finland.

## Author contributions

Conceptualization: S.L. Methodology: E.M., I.T., M.U.T., M.R., N.B., K.H.S., M.K.I., P.C.D. Software: P.C.D. Investigation: E.M., M.R., M.U.T., N.B., I.T., M.G., T.K. Formal analysis: S.L., E.M., P.C.D. Validation: P.C.D., S.L. Resources: A.M.D., M.G., T.K., M.O., P.C.D., A.H. Data curation: E.M., S.L., K.H.S., P.C.D. Visualization: S.L., E.M., P.C.D. Writing – original draft: S.L., E.M. Writing – review & editing: all authors. Supervision: S.L., E.M., A.H., M.K.I. Funding acquisition: S.L., E.M.

## Additional information

**AI use declaration.** The authors used an AI-assisted language tool to proofread the manuscript for grammar, spelling and clarity and to improve conciseness. The scientific content, interpretations, analyses and conclusions, and the original text and ideas, were developed by the authors, who reviewed and approved all AI-assisted edits and take full responsibility for the final content.

## Supplementary figure legends

**Supplementary Fig. S1 | Compound-class composition of the annotated features.** The 514 GNPS2 library-annotated features grouped into compound classes assigned by rule from the library annotation name (class group: fine class). Bars show the number of annotated features per fine class; the text gives the number annotated, the number tested (passing the blank and PEG filters; 285 in total) and the number significantly different in EVs versus the medium in *B. animalis* subsp. *lactis* / *B. longum* 35624 (raw Welch *P* < 0.05 and |log^2^ fold change| ≥ 0.58; exploratory, n = 3 per group). Class assignments and statistics are given in Supplementary Table 3c.

**Supplementary Fig. S2 | Annotated features removed by the blank filter.** Heatmap of the 224 annotated features removed as background by the FBMN-STATS criterion, showing log_2_(peak area + 1) in every sample, including the four solvent blanks; white indicates not detected. Side bars mark candidate PEG annotations (67 features) and features that would have passed the filter if compared with their strongest sample group (top block). Columns are grouped as blanks, medium and the samples of each strain; feature names and values are listed in Supplementary Table 5c.

**Supplementary Fig. S3 | Annotation evidence for vesicle-enriched features. a**, Mirror plot of experimental (top) and library (bottom) MS/MS spectra for the candidate C24 alkamine pentahydroxy bile acid (propagated candidate, Δ*m* −90.0501; MSI level 3). **b**, Abundance (log₂(peak area + 1)) of the putative monohexosyl diacylglycerol (28:0) detected as sodium (top) and ammonium (bottom) adducts across sample groups; both adducts are higher in vesicles than in supernatants, most clearly in *B. longum*. **c**, MS/MS spectrum of the ammonium adduct, showing loss of hexose, water and ammonia to the diacylglycerol fragment (orange) together with the 12:0, 14:0 and 16:0 acyl-glycerol fragments; the diacylglycerol and hexosyl-diacylglycerol features were detected within the *m/z* 700 acquisition window. d, Mirror plot of experimental (top) and library (bottom) MS/MS spectra for nicotinamide (library match, MSI level 2). Symbols and colours as in Fig. 3d; horizontal bars show the group mean; n = 3 biological replicates per group (n = 4 solvent blanks).

**Supplementary Fig. S4 | Volcano plots by compound class, EVs versus uninoculated medium.** The 285 tested annotated features (passing the blank and PEG filters) for *B. animalis* subsp. *lactis* (left) and *B. longum* 35624 (right): log^2^ fold change (pseudocount 1) against −log^10^ raw Welch *P* on log^2^(peak area + 1). Colours and marker shapes give the fine compound class (legend: number tested; number significant in *B. animalis* subsp. *lactis* / *B. longum*); marker size and outline give the *P* value. Significant, raw *P* < 0.05 and |log^2^ fold change| ≥ 0.58 (exploratory, n = 3 per group). Most features are lower in EVs; 31 per strain are higher. The feature with the smallest *P* value in each class group is labelled (GNPS query scan in square brackets); fold changes near ±20 are features with zero area in one group. Statistics are given in Supplementary Table 3c.

**Supplementary Fig. S5 | Non-biological and unannotated features.** a, Features consistent with disinfectants, polymer additives and surfactants. Putative benzalkonium C16 and a tetramethylpiperidinol conjugate (hindered-amine light stabiliser related) are enriched in the vesicle preparations of both strains; putative N-octyl- and N-decyldiethanolamine (whose library matches, the ammonium adducts of dodecanoic and 12-methyltridecanoic acid, are isobaric) are highest in the uninoculated medium and also present in the blanks; a cyclic siloxane is elevated in a single group. b, Unannotated features with culture- or strain-specific patterns: three are produced in the supernatants of both strains, and one is depleted only by *B. longum* 35624. Abundance as log^2^(peak area + 1); symbols and colours as in Fig. 3d; horizontal bars show the group mean; n = 3 biological replicates per group (n = 4 solvent blanks).

**Supplementary Fig. S6 | Global signal shift, shift-corrected class changes and bile acid control features.** a, Per-sample median log^2^ ratio to the mean medium across the 4,648 features detected in all 15 medium and culture samples. b, Median shift-corrected log^2^ change versus the medium per compound class (classes with at least three features; number of features in brackets): only glycerolipids and lysophospholipids exceed two-fold, whereas the N-acyl amino acid and bile acid classes change little and amino acids, di- and tripeptides are not broadly depleted. c, Putative free cholic acid (sodium adduct), a trihydroxy bile acid in-source fragment and a taurine-conjugated dihydroxy bile acid: the free bile acid features rise at 48 h in both strains but with a large spread between replicates, and the conjugate does not decrease. Together these panels indicate that the conjugated-metabolite classes are not produced de novo by either strain under these conditions. Values are given in Supplementary Table 2b,c.

**Supplementary Fig. S7 | Strain-specific conversions in the soluble exometabolome.** Abundance (log₂(peak area + 1)) in the uninoculated medium, the 24 h and 48 h supernatants and the EVs of *B. animalis* subsp. *lactis* and *B. longum* 35624, and in solvent blanks. **a**, Tryptophan catabolism: putative indole-3-lactic acid appears essentially only in *B. longum* and rises from 24 to 48 h, whereas tryptophan is unchanged. **b**, Purine metabolism: *B. animalis* subsp. *lactis* depletes adenosine, guanosine and adenine and accumulates hypoxanthine, with the opposite pattern in *B. longum*. **c**, Carbohydrate use: a putative trisaccharide and a putative free hexose are cleared or strongly depleted by *B. longum* but not by *B. animalis* subsp. *lactis*, whereas a putative disaccharide is unchanged in both. Symbols and colours as in Fig. 3d; horizontal bars show the group mean; n = 3 biological replicates per group (n = 4 solvent blanks); comparisons are exploratory and test statistics are given in Supplementary Table 2.

**Supplementary Fig. S8 | Interpretive schematic of the strain-specific conversions.** a, Nicotinamide deamidation and b, indole-3-lactic acid formation in *B. longum* 35624; c, purine handling in *B. animalis* subsp. *lactis*; d, carbohydrate use in *B. longum* 35624. Solid boxes, observed; dashed boxes, inferred intermediates or products not measured here; enzyme names on arrows are inferred.

## Supplementary Methods

### Feature finding

Everything Bagel (version 2.0.0+v2) was used for feature finding because it processes the complete dataset rapidly and in a fully automated way on GNPS2. Because it is a recently introduced workflow, its output was interpreted with care, particularly for low-abundance features close to the adaptive noise threshold, and all downstream filtering, annotation and statistical steps were performed in documented R scripts. Library search parameters: [minimum cosine, minimum matched peaks, precursor and fragment tolerances to be added from the GNPS2 job].

### Linking library annotations to features

GNPS2 library matches were linked to features through the GNPS query scan number. Because overlap counts cannot distinguish between candidate identifiers, the link was verified for each dataset by joining each candidate identifier (query scan and query index) to the feature table and confirming that the precursor *m/z* of the library match agreed with that of the feature (±0.02 Da); the identifier with the highest agreement was used. Where several library matches existed for a feature, the match with the highest cosine score was retained, and the fate of every library match (used, or removed and why) was recorded.

### Documentation of blank filtering and contaminant exclusion

For every feature, the mean intensity in the blanks and in each sample group was recorded, together with the ratio used for filtering and whether the feature would have passed if compared with its strongest sample group rather than with the mean of all samples (Supplementary Table 5; Supplementary Fig. S2). Candidate PEG annotations were identified from their library names, and their detection in blanks and in each sample group was tabulated separately. PCA including the blanks was computed on all features before and after blank filtering to visualise the effect of the filter. For display, adduct, collision-energy and library metadata were removed from compound names; full names are given in the supplementary tables.

